# BGMDB: A curated database linking gut microbiota dysbiosis to brain disorders

**DOI:** 10.1101/2024.06.21.599994

**Authors:** Kai Shi, Pengyang Zhao, lin Li, Qiaohui Liu, Zhengxia Wu, Qisheng He, Juehua Yu

## Abstract

The gut microbiota plays a pivotal role in human health by modulating physiological homeostasis and influencing the pathogenesis of various diseases. Recent studies have underscored the close relationship between neurotransmitters, which act as communication mediators between the gut and brain, and the development and treatment of multiple brain disorders. Despite these advances, the intricate interactions between gut microbiota and brain diseases remain largely unexplored in the extensive biomedical literature. There is a notable absence of a structured database focusing on gut microbiota-brain disease associations. Introducing BGMDB (Brain Disease Gut Microbiota Database), a meticulously curated database designed to provide experimentally supported connections between gut microbiota and brain diseases. The current version of BGMDB extensively covers 1,419 associations involving 609 gut microbiota and 43 brain diseases, including 184 specific association triplets linking brain diseases, neurotransmitters, and gut microbiota among six neurotransmitters. Noteworthy is that BGMDB integrates gene data related to gut microbiota from the gutMGene database. Brain region and disease microbial networks are introduced to investigate potential common genetic relationships between brain diseases and brain region changes. Each entry in BGMDB offers detailed insights into specific associations, including the particular brain disease implicated, the involved gut microbiota, neurotransmitter, and a concise description of the relationship supported by relevant literature references. To facilitate easier access to relevant information for specific brain diseases, BGMDB provides enhanced graphical query options to address various biologically pertinent inquiries. Additionally, a user-friendly interface allows users to browse, retrieve, and download entries conveniently. BGMDB serves as a valuable resource for investigating microbes associated with human brain disorders. Access BGMDB through http://43.139.38.118:8080/demo02/.

## Introduction

The human gut microecosystem harbors an extensive array of microbes, including up to 38 trillion bacteria, eukaryotes, viruses, and archaea, predominantly residing in the intestines^1, 2^.. As the largest and most intricate ecological community within the human body, the gut microbiota and its metabolites play a critical role in maintaining physiological homeostasis and influencing disease onset, thereby being indispensable for human health^3, 4^. Recent evidence strongly suggests that alterations in the diversity and composition of gut microbes are closely linked to the pathogenesis of numerous brain diseases. For instance, studies have consistently demonstrated that Alzheimer’s disease, Parkinson’s disease, and amyotrophic lateral sclerosis exhibit indications of intestinal microbial dysfunction^5^. The microbiota provides hosts with genetic, transcriptomic, and metabolic attributes that impact host biology both beneficially and detrimentally. Over recent decades, it has become increasingly evident that microbial communities are crucial for human and environmental well-being. Serving as information transmitters in the nervous system, neurotransmitters play a significant role in nervous system function. Recent research has emphasized the close association between neurotransmitters and the development and treatment of a wide range of brain diseases^6^.

With the rapid advancement of high-throughput metagenomic sequencing technologies, an increasing number of connections have been discovered between gut microbiota-disease relationships and neurotransmitters^7^. These publicly available resources serve as vital references for studying disease pathogenesis. However, despite the growing body of related studies, users still face challenges in accessing, processing, and analyzing data as research on the associations between diseases and microbiota remains relatively scattered. In response, Janssens et al. introduced Disbiome^8^, a standardized database that collects and presents published information on microbiota and diseases. Additionally, Oliveira et al. developed MicrobiomeDB^9^, a platform for data discovery and analysis that empowers researchers to effectively query microbiota and disease datasets using experimental variables. Subsequently, Zeng et al. created MASI^10^, which enables retrieval and browsing of correlation information between gut microbiota and host metabolites. It also offers visualization of correlation network graphs and scatter plots with a specific focus on the association between gut microbiota and host metabolites. Overall, while various databases store gut microbiota and disease data, including some neurological diseases, none provide a comprehensive integration of gut microbiota changes with neurological or brain diseases. Worth noting is the introduction of BrainBase^11^ by Liu et al., which provides knowledge on brain disease-gene associations and drug-gene interactions. It identifies brain-specific genes, mines characteristic genes of glioma, and offers visualization maps of multi-omics data, thus providing essential data resources for understanding the pathogenesis and development mechanisms of brain diseases. Unfortunately, Liu et al. did not explore the association between brain diseases and gut microbiota. Sun et al. recently constructed and developed MMiKG^12^, a data platform based on the knowledge graph construction of the psychiatric disease-intermediate-gut microbiome association. However, the main purpose of the MMiKG platform is to predict hidden relationships between nodes, and it focuses solely on psychiatric disorders.

In response to these gaps, we have developed BGMDB, a meticulously curated database that documents experimentally supported associations between gut microbiota, brain diseases, and neurotransmitters, offering a user-friendly visual interface. The current version of BGMDB includes 1,419 association entries between 609 microbiota and 43 brain diseases. It also integrates related neurotransmitter information from the MiKG4MD^13^ database and gene data related to microbiota from the gutMGene^14^ database. Standardization of terms for brain diseases and gut microbiota was based on the National Center for Biotechnology Information (NCBI) taxonomy and Medical Subject Headings (MeSH) disease categories. Additionally, the BGMDB database offers data retrieval and download functions, enabling users to access and download necessary data for their research and analysis as per their requirements. Our aim is for BGMDB to serve as a valuable resource for investigating the relationship between human gut microbiota and brain diseases, thereby enhancing our understanding of the link between gut microbiota and overall brain health.

## Methods

BGMDB is a database containing experimentally supported microbe-brain disease associations. Through an analysis of thousands of studies, we have identified and documented 1419 associations between 609 microbiota, 43 brain diseases, and 6 neurotransmitters (Table 1).

**Table 1.** Microbial species AND Brain disease data in the database.

|  | Classes | Amount | Total |
| --- | --- | --- | --- |
| Types of brain diseases | Mental Disorders | 17 | 43 |
|  | Nervous System Diseases | 26 |  |
| Microbial Species | Kingdom | 2 | 609 |
|  | Phylum | 18 |  |
|  | Class | 16 |  |
|  | Order | 25 |  |
|  | Family | 71 |  |
|  | Genus | 206 |  |
|  | Species | 122 |  |
|  | Other | 149 |  |
| Neurotransmitter | Cholines | 1 | 6 |
|  | Monoamines | 4 |  |
|  | Other | 1 |  |
| Encephalic Region | Brodmann Areas | 8 | 8 |

## Data collection and processing

### Collection of raw data

Our primary objective is to provide comprehensive and accurate information regarding experimentally supported associations between gut microbiota and brain diseases. To ensure the quality of our data, we manually extracted associations from various sources, including PubMed^15^, Web of Science, and other relevant databases such as MicroPhenoDB, Peryton^16^, and gutMDisorder^17^. By conducting keyword searches with terms like “microbiota,” “microbiome,” “microbe,” “bacteria,” “gut,” “intestinal,” “brain disease,” “brain disorder,” “gut microbiota,” “gut flora,” “intestinal bacteria,” “neurotransmitter,” “serotonin,” “dopamine,” “norepinephrine,” “GABA,” “histamine,” and “acetylcholine,” we identified a total of 1,445 research articles. From these articles, we manually extracted interaction records reported in each publication. Whenever available, we also collected experimental details such as sample source, sequencing method, processing software, etc., pertaining to the interactions between brain diseases and gut microbiota. Additionally, we incorporated neurotransmitter correlations from the MiKG4MD database into our version. Furthermore, information including gene name, gene ID, and sequence was obtained from the gutMGene database. Lastly, we conducted a thorough manual assessment of the literature to enhance the reliability of the associations between gut microbiota and brain diseases.

### Data processing

The latest iteration of BGMDB comprehensively catalogs extensive data on brain diseases and gut microbiota. Given the diverse descriptions of microbiotas and diseases in the literature, the standardization of entity naming is essential to enhance database accessibility. To achieve this, we utilize MeSH for standardizing disease names and unify microbiota names using the NCBI database. Moreover, the gathered literature and experimental data undergo filtration and cleaning processes to eliminate duplicate and uncertain information, ensuring data quality and reliability.Database content and usage

## Database contents and Usage

### Database Contents

BGMDB is a comprehensive database dedicated to exploring the association between human brain diseases and gut microbiota. It serves as a valuable resource for the study of human health and disease. The current version of BGMDB contains 1,419 associations between 609 gut microbiota and 43 human brain diseases. Each brain disease entity within the database encompasses various fields, including disease name, related references, experimental data, and gut microbiota associations. Similarly, the microbiota entity comprises microbiota name, classification information, references, and related disease information. The related references provide details such as title, author, journal, and abstract, while the experimental data encompass experiment type, design, sample size, and more. Furthermore, we have gathered and organized information on neurotransmitters that interact with specific brain diseases and gut microbiota. Additionally, gene information of gut microbiota, including gene name, gene ID, and sequence information, has been recorded. Users can conveniently access the desired information by entering keywords or selecting specific categories of disease, microbiota, or neurotransmitter. Moreover, relevant data can be downloaded from the “Download” interface in CSV format, enabling users to conduct in-depth analysis (Figure 1).

**Figure 1.**
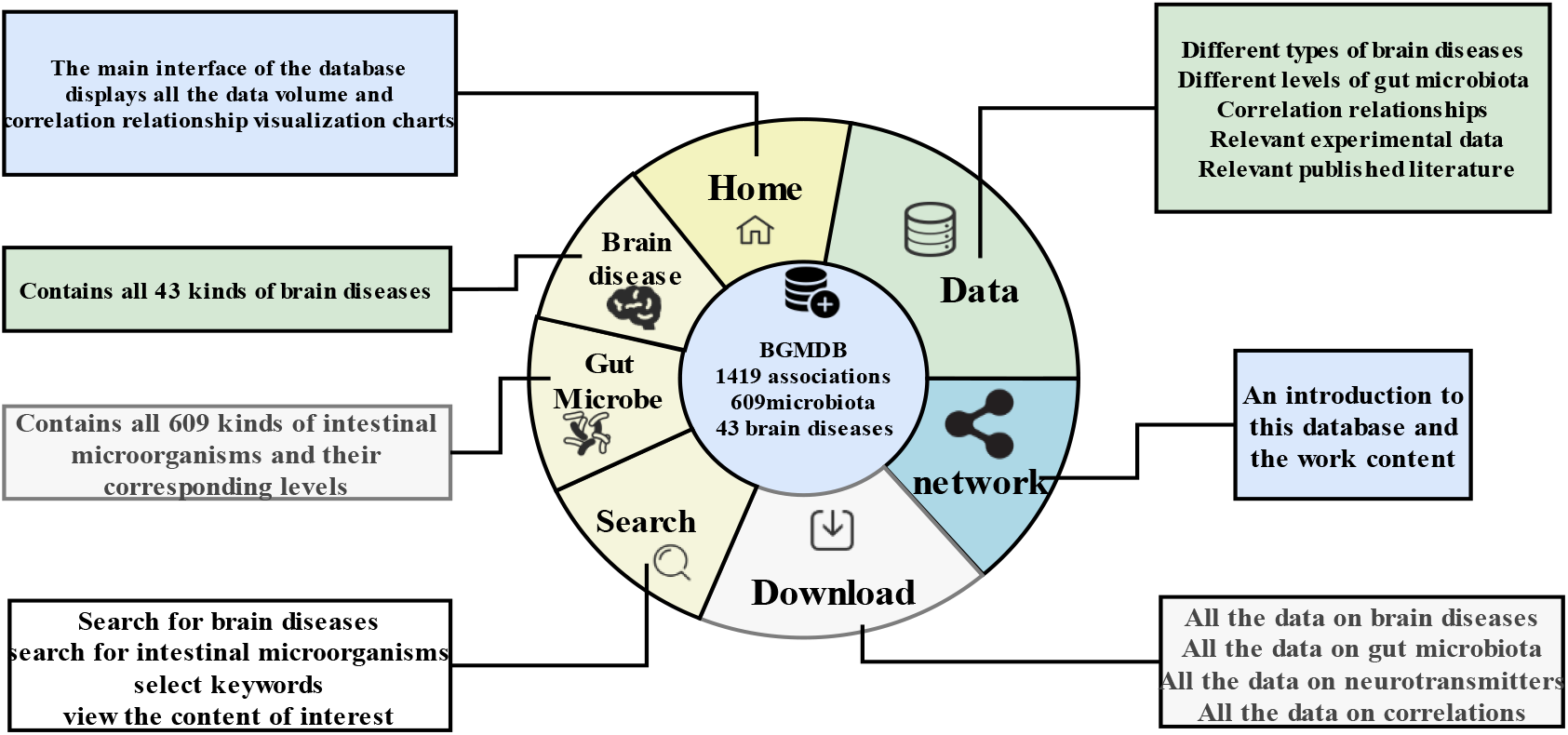
BGMD Function Display and Search. The search function allows users to retrieve and view relevant data. The data interface provides a detailed view of the data, with the option to click on specific data points for more information in a pop-up window. The Data-Download interface

### Encephalic Region and Disease-Microbe Network

The term encephalic region (ER) is derived from medical studies on various functional areas of the brain, combined with theories from modern scientific psychology^18^. The most commonly used method to divide the brain into distinct regions is the Brodmann Areas, which categorize the brain into 52 regions. The brain acts as the central control center for human behavioral activities, often requiring coordinated efforts from multiple brain regions. Zhao et al. ^19^ found that abnormalities in brain structure can lead to the occurrence of a variety of diseases, and there may be a common genetic relationship between brain diseases and changes in brain regions.

Based on this hypothesis, we conducted research to explore the correlation between brain diseases and specific brain regions while also examining their relationship with intestinal microbiota. Within the interface of brain regions, users have the option to select specific regions to access comprehensive data on diseases associated with each region and vice versa. However, due to limitations in the available data, this database currently includes association data between eight encephalic regions and 12 types of brain diseases (Figure 2A).

**Figure 2.**
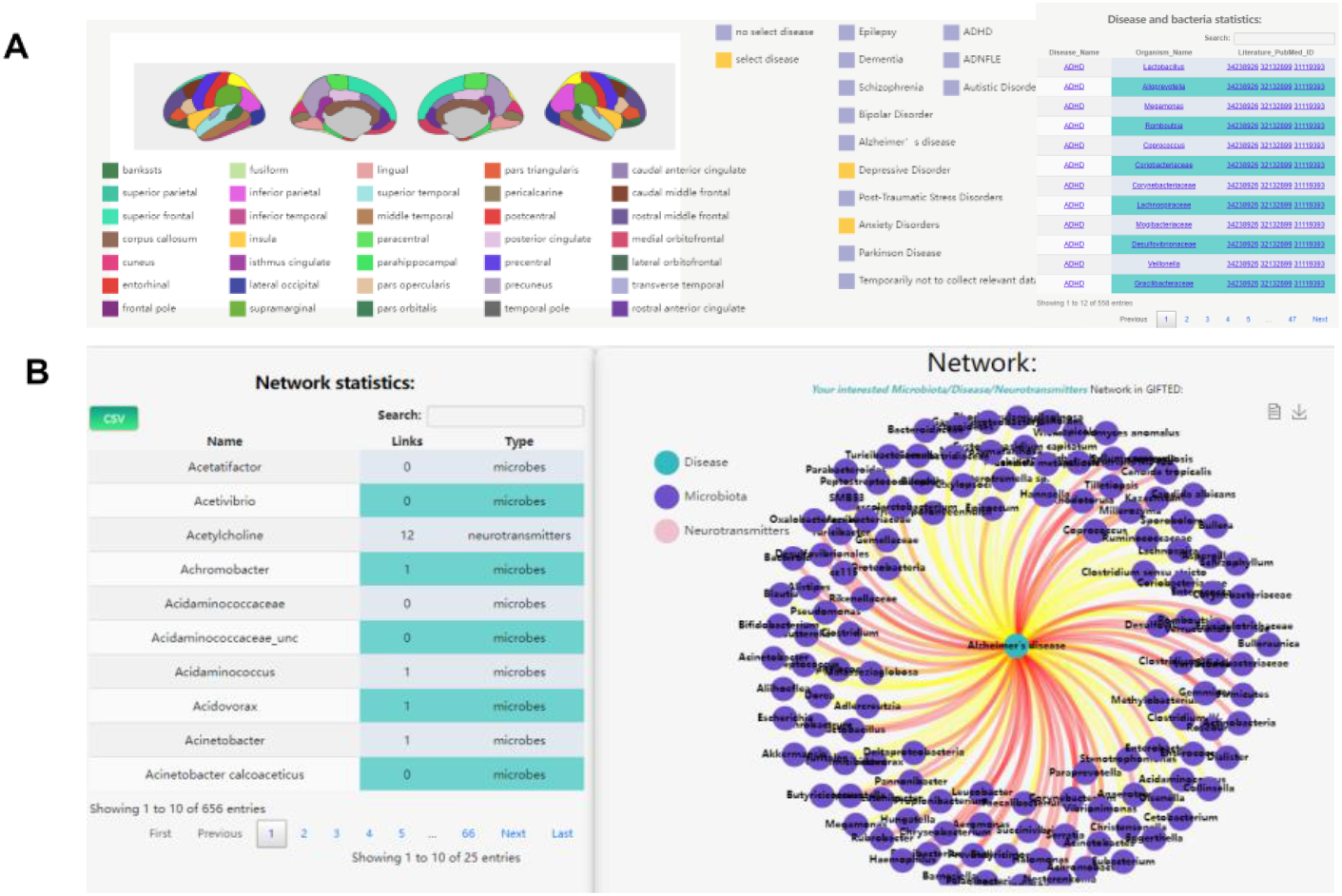
The Encephalic Region and Network Function Schematic Diagram.

To further investigate the relationship between brain diseases and intestinal microbiota, we have introduced the Brain Disease-Intestinal Microbe Relationship Network. Users have the ability to select specific diseases or microbes of interest to view the associated data. These associations are presented in the form of networks and tables, providing users with clear and visual representations of the relationships (Figure 2B).

### Database design

The web application of BGMDB is developed using the Java language and constructed on the model-view-controller model and Spring Boot framework. It is deployed on the Apache Tomcat web server. Data access, search functionality, and visualization features are implemented utilizing Ajax API technology. The browsing and search web interface is created using Java Server Pages (JSP). This architecture facilitates the provision of a comprehensive data resource for researchers (Figure 3).

**Figure 3.**
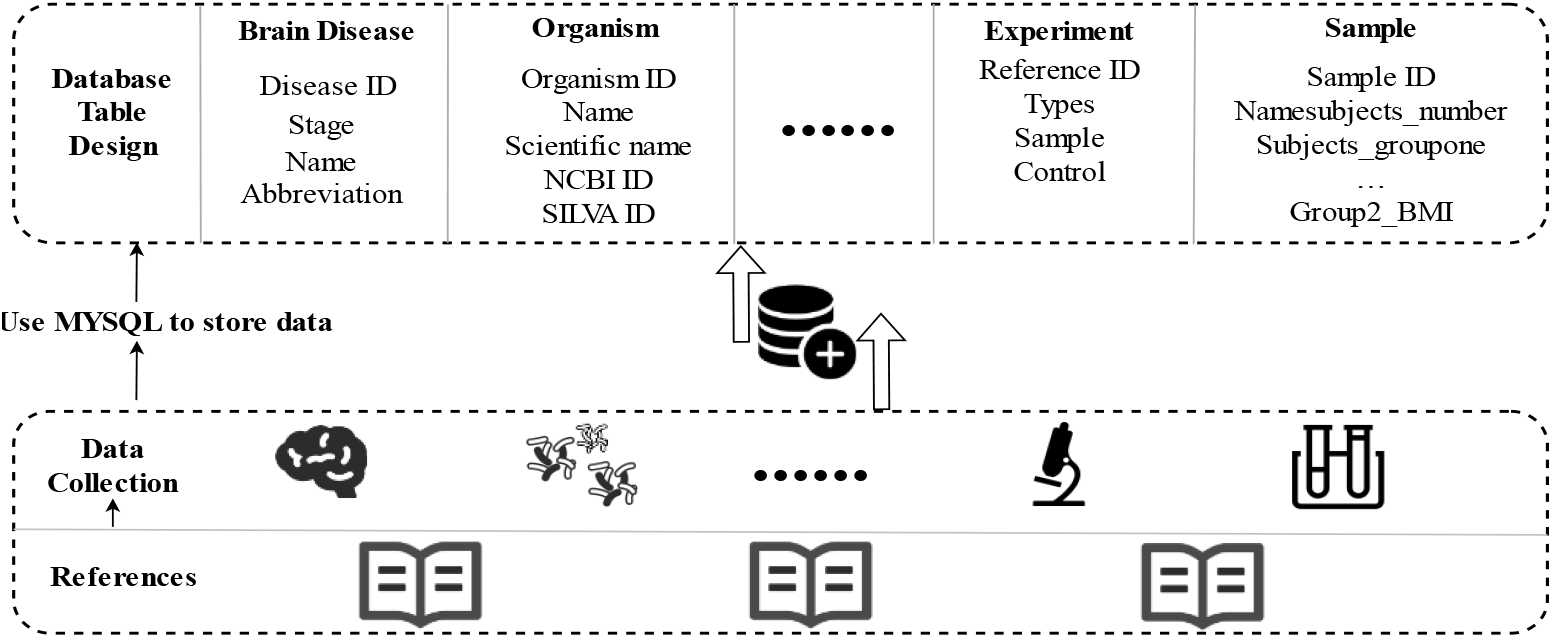
BGMD Construction Process. To construct the BGMDB, a comprehensive collection of data was meticulously gathered from a diverse range of carefully selected literature sources. This data was then organized, modified, and subsequently used to create data tables and databases specifically tailored for the BGMDB.

**Figure 4.**
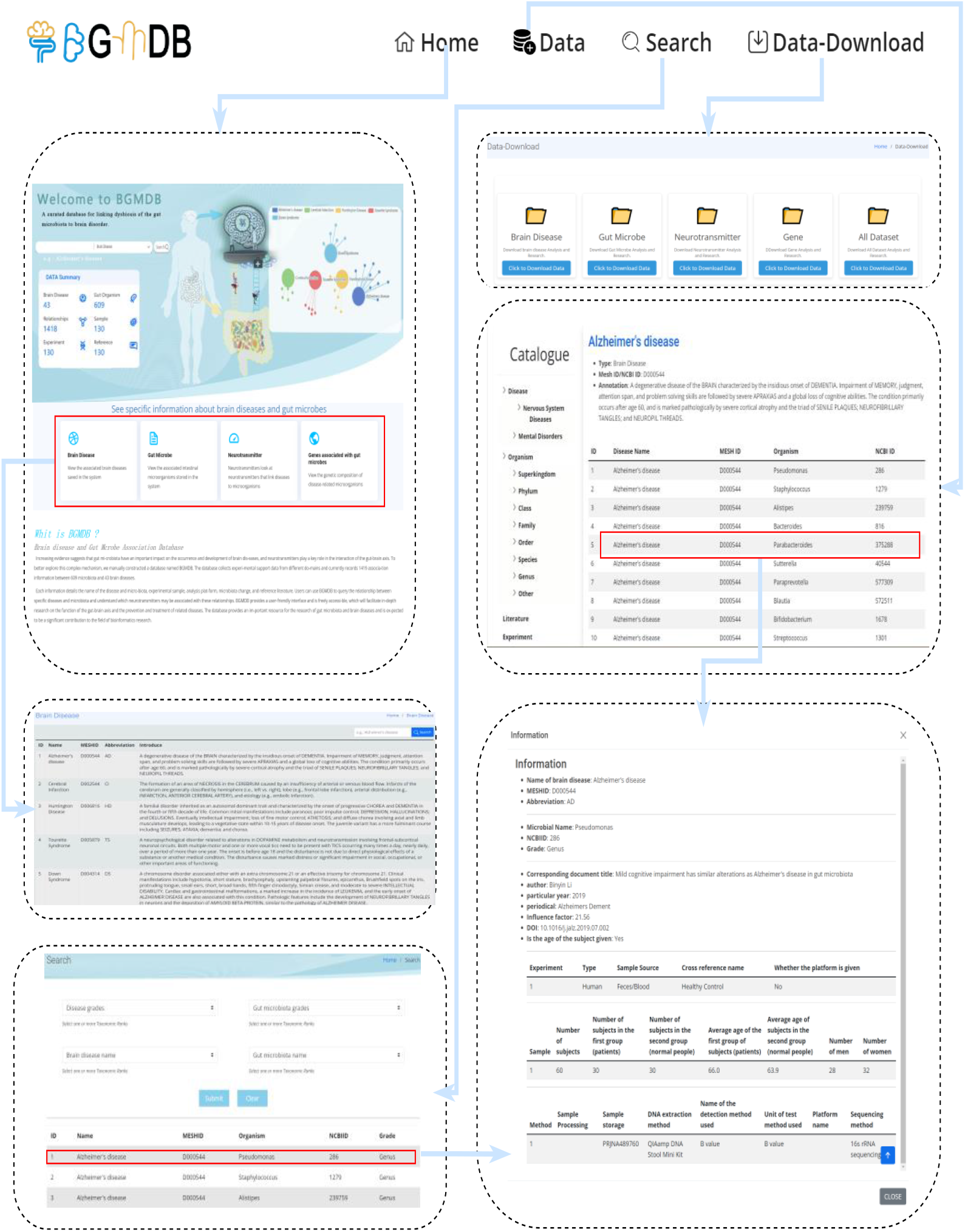
BGMDB Function Page

## Results

### Web interface

The platform offers a user-friendly web interface that allows users to search, browse, download, and analyze the associations between gut microbiota and brain diseases. Furthermore, the website includes specialized search applications for specific microbiota or diseases, enabling users to access prioritized relationships. Users can also download the prioritized associations of microbiota and diseases as CSV files for further analysis. The hierarchies of microbiota and diseases are individually presented on the “Data” webpage. To ensure robust performance, we conducted testing of the BGMDB website across various web browsers, including Mozilla Firefox, Google Chrome, and Internet Explorer.

## Usage Notes

Recent studies have highlighted a significant relationship between gut microbiota and brain diseases such as Anxiety^20^, Depression^21^, and Parkinson’s disease^22^. Researchers leverage data from various public databases to delve into this correlation. BGMDB stands out as an online knowledge-based database offering the most up-to-date insights into the connection between gut microbiota and brain disease. By clicking on the name of a specific brain disease or gut microbiota, users can access detailed information, including the MeSH ID and abbreviation of the brain disease, along with related experiments, samples, associated gut microbiota, and relevant references.

Researchers utilize the data within BGMDB to scrutinize the alterations in gut microbiota communities among patients with brain diseases and investigate the link between these changes and disease progression. To strengthen comprehension of BGMDB’s utilization, we present application cases.

### Case 1: Mining ho ologo s rain Diseases

Homologous brain disorders share a common etiology and symptoms, necessitating personalized treatment approaches. Concurrent treatment of such homologous brain diseases can significantly improve therapeutic outcomes, reduce the treatment burden, refine disease management, and lower social costs. Our goal is to pinpoint distinct disorders that are associated with analogous intestinal microbiota within the gut-brain axis knowledge. For example, researchers can navigate through the “Data” and “Retrieval” interfaces to discover that Alzheimer’s disease is classified under Central Nervous System Diseases. Similarly^23^,, Tourette Syndrome, Attention Deficit Disorder with Hyperactivity^24^, and Autistic Disorder^25^, among others, are considered parallel brain disorders. Furthermore, Neurotic Anorexia, Anxiety Disorder, Bipolar Disorder, and others fall within the same category as Central Nervous System Diseases (Figure 5).

**Figure 5.**
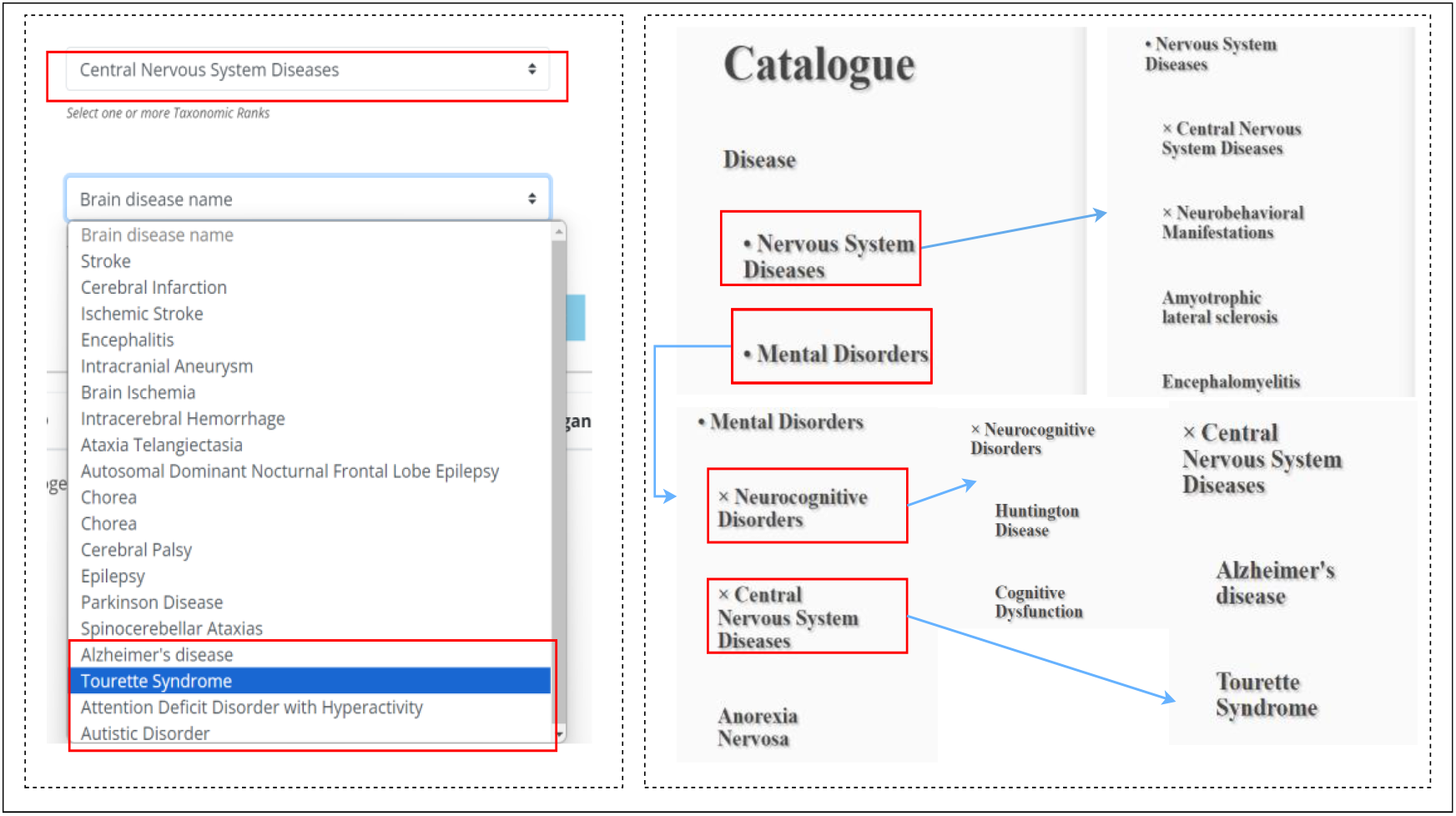
Mining homologous Brain Diseases.

### Case 2: Mining biotic Microbes

Currently, the treatment landscape for brain disorders heavily relies on pharmaceuticals and psychotherapy, yet their effectiveness remains limited. Exploring the intricacies of the brain-gut axis reveals new opportunities for managing brain disorders. By examining these microbial entities, valuable insights for drug development could be uncovered. Understanding the interactions and common pathways among microbes can reveal new targets and therapeutic approaches. In this context, researchers can access hierarchical classifications of identical microbes and associated disease types through the “Retrieval,” “Data,” and “Network” interfaces. They can also explore the gut microbiota correlated with Alzheimer’s disease and its taxonomic levels, such as Coriobacteriaceae^26^ at the family level, Staphylococcus^27^ at the genus level, and others. Furthermore, researchers may note that both stroke disease patients and Alzheimer’s disease patients harbor Pseudomonas data readily accessible through the “Data-Download” section (Figure 6).

**Figure 6.**
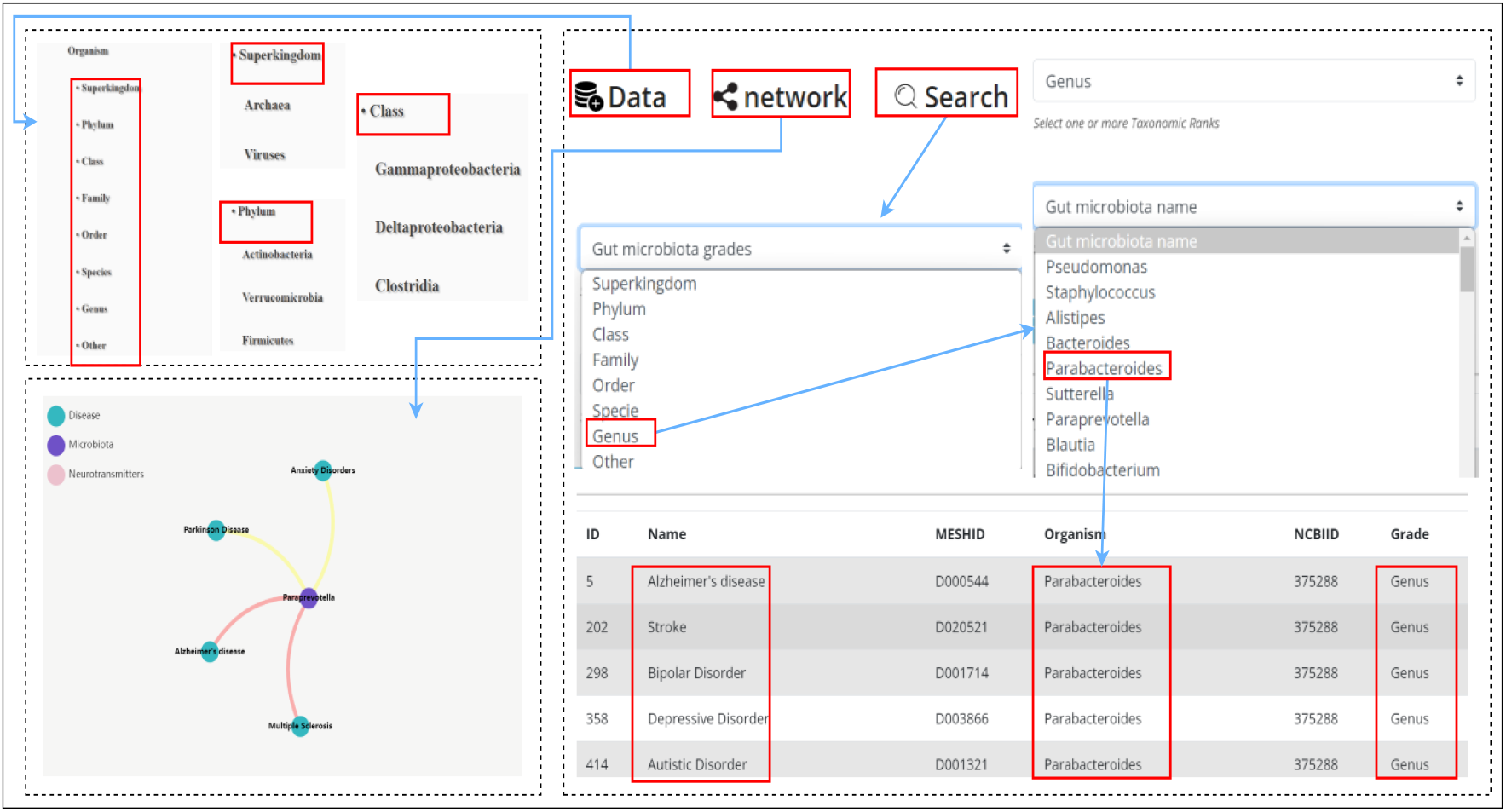
Mining symbiotic microorganisms.

### Case 3: In estigating Encephalic Region-Disease Relationships and the Role of Ne rotrans itters in rain Disease-t Microbiota ssociations

The brain serves as the central hub for regulating human behavior, requiring coordinated efforts across multiple brain regions for every activity. Identifying the specific regions within the brain where diseases originate can greatly boost targeted therapeutic effectiveness and provide diverse treatment approaches for brain disorders. Our database provides insights into the associations between brain regions, diseases, and gut microbiota. For example, researchers can click on the Superior Parietal region, and BGMDB will display associated brain diseases such as Anxiety and Depressive disorders along with the corresponding gut microbiota on the right side (Figure 7).

**Figure 7.**
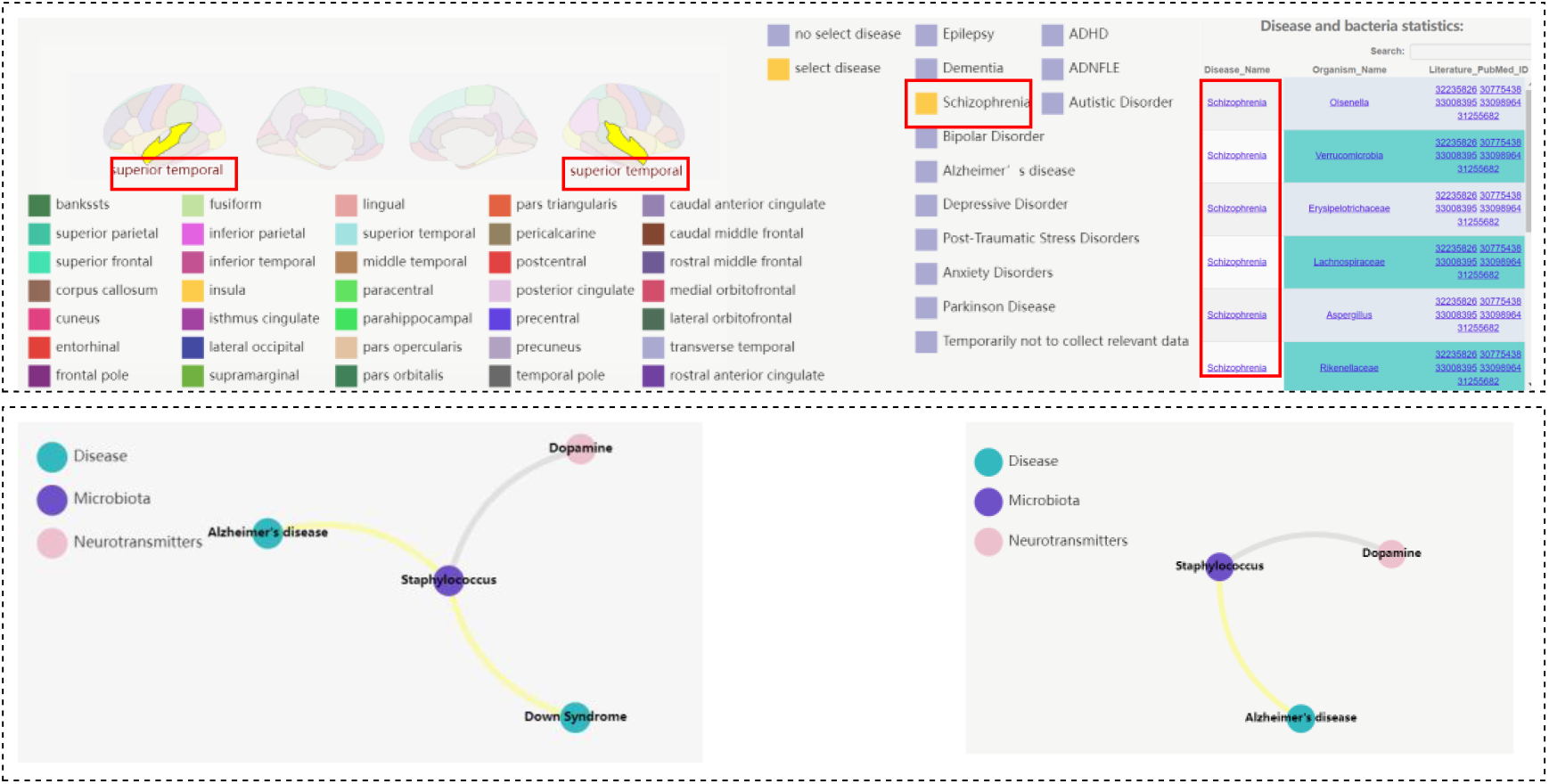
Mining brain regions and neurotransmitters.

Neurotransmitters play a crucial role as intermediate messengers within the brain-gut axis, attracting significant interest from researchers due to their involvement in the interplay between microbes and brain disorders, which is pivotal for diagnosis and treatment. In the “Network” section of BGMDB, researchers can explore the relationship network between brain diseases, gut microbiota, and neurotransmitters. This allows them to understand, for instance, how variations in Lactobacillus plantarum bacteria in Alzheimer’s disease patients are influenced by Acetylcholine^28^ and how variations in Escherichia bacteria in Alzheimer’s disease patients are modulated by Serotonin^29^.

With the information and insights gained from these findings, researchers can utilize the data from BGMDB to pinpoint potential targets and drug candidates. Comparative analysis within BGMDB enables exploration of differences in gut microbiota between patients with various brain disorders and healthy individuals as well as potential synergistic or antagonistic effects. BGMDB proves to be an indispensable resource for researchers, offering novel perspectives and methodologies in the field of bioinformatics.

## COMPARISON WITH EXISTING DATABASE ARCHITECTURE RESOURCES

Online resources such as Disbiome, gutMDisorder, Peryton, and MASI provide similar content to BGMDB, focusing on the associations between microbiota and diseases. However, they have different intentions and purposes. The Disbiome database primarily focuses on changes in microbiota composition across various diseases. The gutMDisorder database emphasizes the effects of intervention factors like drugs and food on gut microbiota dysbiosis. The Peryton database examines the relationship between gut microbiota and host phenotypes in animal models. The MASI database specifically investigates the association between gut microbiota and host metabolic products. In contrast, the BGMDB database encompasses a broader range of associations, particularly related to brain disease health phenotypes. It hosts associations between different types of brain disease phenotypes and other factors. These entries significantly advance the understanding of gut microbiota-brain disease associations, offering a systematic collection that can be utilized to study the potential connections between gut microbiota and brain diseases for the first time. In addition to its main content, BGMDB provides detailed association information between specific brain diseases and gut microbiota. This includes specific neurotransmitter information related to brain diseases and gene data related to gut microbiota. The database classifies gut microbiota based on hierarchical levels such as “kingdom,” “phylum,” “class,” “order,” “family,” “genus,” and “species.” Diseases are also categorized into two main groups: neurological diseases and mental disorders. BGMDB supports advanced queries using one or more brain disease and gut microbiota names as well as multiple taxonomic ranks. Additional query options include filtering by “disease,” “experimental method,” “sample source,” “sample size,” and associations derived from healthy controls only^16^. Further distinguishing BGMDB from existing resources are several fine annotation and curation choices. These include highlighting information on commonly studied brain diseases and gut microbiota in query results, adopting a systematic and unified nomenclature for brain disease names and gut microbiota, and implementing dedicated associations specifically for brain diseases and gut microbiota.

## DISCUSSION AND FUTURE WORK

The gut microbiota plays a crucial role in disease development and human health, impacting conditions such as intestinal diseases, neurologic diseases, and even several cancers^30-32^. Over the years, researchers and clinicians have sought to uncover the link between intestinal microbiota and brain diseases in order to improve diagnosis and treatment strategies. It is believed that some brain diseases are not solely genetic but also microbial in nature. Advancements in high-throughput technologies have led to a significant increase in experiments exploring associations between gut microbiota and brain diseases. As a result, a wealth of experimentally supported associations are scattered across thousands of published studies. Extracting and organizing these associations from the vast literature landscape and constructing association databases play a crucial role in furthering our understanding of the relationships between gut microbiota and brain diseases.

To create a comprehensive repository on the roles of microbes in brain diseases, we have introduced BGMDB, a meticulously curated database that catalogs associations between gut microbiota and brain diseases^33^. This database compiles 1,419 associations involving 609 gut microbiota and 43 human brain diseases. The primary aim is to elucidate the intricate relationship between gut microbiota and brain diseases, leverage the wealth of accumulated data from recent years, and aid researchers in uncovering novel insights. Drawing data from over 1,400 articles, our database not only consolidates information but also conducts thorough analyses and constructs visual networks. By serving as a valuable and user-friendly tool, researchers can extract important implications from this extensive interconnected resource.

In order to optimize data collection quality and streamline the process, we propose implementing the following three strategies. Firstly, we will employ text-mining tools to filter brain disease-related literature by analyzing titles and abstracts. Secondly, regular reviews will be conducted on existing database entries to ensure data accuracy. Finally, updates on gut microbiota and brain disease data will be made quarterly to incorporate the latest findings.

A recent study by LR et al.^34^ introduced a novel approach to transform microbiota data and networks into graph databases. This method leverages the efficiency, flexibility, and scalability of graph databases and includes the development of the mako tool. The mako tool enables users to explore patterns and relationships within microbiota data and networks, including microbe distribution, composition, function, interactions, as well as associations with host phenotypes, disease status, environmental factors, and more. The introduction of mako offers fresh perspectives and methodologies for microbiota research. It also presents new possibilities for the future direction and content of the BGMDB database, specifically in terms of transforming microbiota data and networks into graph databases and visualizing query results. This will allow users to perform network-based queries effectively. This facilitates user analysis of microbiota data and networks in a rapid, adaptable, user-friendly, and scalable manner. As high-throughput technologies continue to advance, more characteristics of gut microbiota will be unveiled, potentially leading to a significant increase in microbial resources. Therefore, it would be highly beneficial to integrate BGMDB with other relevant resources to offer a convenient means of annotating the functions of gut microbiota. In pursuit of this goal, further connections will be established between gut microbiota, brain diseases, neurotransmitters, and other potential related resources^35^. In addition to the ongoing collection of new human gut microbiota data in the years ahead, our plan includes expanding the content of BGMDB. This expansion will encompass, but not be limited to, in-depth exploration of the relationship between gut microbiota and brain diseases by integrating genetic and environmental factors; comprehensive investigation of the association between gut microbiota and brain diseases by incorporating other relevant factors such as genetic and environmental factors; and the addition of data analysis tools based on artificial intelligence technologies such as machine learning and deep learning^36, 37^. These enhancements will further facilitate research on the brain-gut axis and contribute to a deeper understanding of the connections between gut microbiota dysbiosis and brain diseases.

## Acknowledgements

The authors thank the anonymous referees for suggestions that helped improve the paper substantially. Research supported by the National Natural Science Foundation of China (62162019, 62166014), Shanghai Municipal Science and Technology Major Project (No. 2018SHZDZX01), Key Laboratory of Computational Neuroscience and Brain-Inspired Intelligence (LCNBI), and ZJLab, Guangxi Key Laboratory Fund of Embedded Technology and Intelligent System, and the startup Grant in Guilin University of Technology. The authors declare that they have no conflicts of interest.

## Data Availability

BGMDB serves as a valuable resource for investigating microbes associated with human brain disorders. Access BGMDB through http://43.139.38.118:8080/demo02/Index/download.

## Code Availability

All software and pipelines were executed according to the manual and protocols of the published bioinformatics tools. The version and code/parameters of software have been described in Methods.

## Author contributions

Kai Shi and Pengyang Zhao carried out the study and drafed the manuscript; Pengyang Zhao, lin Li, Qiaohui Liu, Zhengxia Wu and Qisheng He participated the data processing; Kai Shi directed the project, discussed and revised the manuscript.

